# Operant Behavior in Model Systems

**DOI:** 10.1101/058719

**Authors:** Bjöern Brembs

## Abstract

In contrast to the long-held assumption that the organization of behavior is best characterized as the perception of a sensory stimulus followed by appropriate response (i.e., “sensorimotor hypothesis”), recent converging evidence from multiple systems and fields of study instead suggests that both ancestral and extant general brain function is best described in operant terms. Rather than specifyng precise behaviors, sensory information - if at all present - interacts with ongoing neural activity to instruct the organism which type of spontaneous, exploratory behavior to generate. Evaluating the ensuing reafferent feedback modifies the nervous system such that ongoing neural activity patterns become biased towards activity that has generated increased appetitive and decreased aversive feedback in the past. The neurobiological mechanisms underlying both the exploratory, spontaneous behaviors as well as those underlying the modifications caused by the feedback are becoming increasingly understood, even on a molecular level. It is straightforward to hypothesize that the constant interaction between ongoing neural activity and the incoming sensory stream allows the organism to balance behavioral flexibility with efficiency to accomplish adaptive behavioral choice in an often unpredictably changing environment.

## 1 Operant and Classical Conditioning

One of the traditional dichotomies in learning and memory research is that between operant (instrumental) and classical (Pavlovian) conditioning. The distinction between operant and classical conditioning is merely operational:these terms denote *how* a learning experiment was conducted. In classical conditioning, the animal’ s behavior has no influence over the stimuli it is presented with, while in operant conditioning the animal is in control at least of the most relevant stimuli. The stimuli and behaviors in both classes of experiments can be almost arbitrarily exchanged, as long as these rules are obeyed. Already in the 1930s, Skinner and his contemporaries realized that, in its simplest form, one could conceptualize *what* is being learned in a classical conditioning experiment in only one single process, namely the association between two stimuli. Following Pavlov, this concept can be described as “ stimulus substitution” where the conditioned stimulus (CS; e.g. a tone) receives some of the response-eliciting properties of the unconditioned stimulus (US; e.g. food). Note that this does not preclude the design of less minimal experiments where the animals are allowed to learn more. Operant conditioning, on the other hand, most likely comprises at least two, if not more processes:the association between two stimuli and the association between the behavior of the animal and one or more stimuli (Skinner, 1935, Skinner 1937; Konorski and Miller, 1937a, Konorski and Miller 1937b). This is what Skinner called the “three-term-contingency” of (I) the stimuli that are present just before and during the behavior, (II) the behavior itself and (III) the stimuli that the animal is presented with as a consequence of its behavior (Skinner, 1969). Phrased differently, it was Pavlov’s strike of genius to prevent his dogs from interfering with the experiment by firmly tying them down, as it significantly reduced the number of possible learning processes to those involved in processing external stimuli.

From this brief recapitulation it becomes clear that already 80 years ago, there was very little intrinsic reason to specifically juxtapose these two types of experiments - the reason these classes of experiments are still taught and discussed together today (e.g., Domjan, 2016) are more historical than logical. In these eight decades, a wealth of literature has accumulated describing experiments in which seemingly every possible relation and detail in the configurations of stimuli and behaviors has been tested to compare and contrast operant and classical conditioning. Despite all these efforts, until the advent of modern neurobiological techniques, the more fundamental questions that Skinner and his contemporaries started tossing around in the 1930s remained largely unanswered.

### 1.1 Spontaneous Actions

While the relationship of the animal’s behavior to antecedents was always intuitively obvious in classical conditioning - after all, the US was chosen such as to elicit a clear response - this relationship remained hotly debated in operant conditioning until rather recently. Despite emphasizing the exploratory or spontaneous nature of the (‘emitted’, not ‘elicited’) behavior in operant conditioning experiments (Skinner, 1938), the behavior was nevertheless routinely referred to as a ‘response’, implying some eliciting factor (Dickinson, 1985). This ambiguity about whether or not one needed to describe the behavior of animals in classical and operant conditioning experiments in different terms has provoked discussions until this day (Domjan, 2016). Arguably, the ambiguity may be traced to the behaviorist effort to separate behaviorism from the prevailing animist psychology of the early 20^th^ century on one side, while at the same time emphasizing the difference to the more conventional schools of classical conditioning, e.g., reflexology, on the other. As demonstrating spontaneous behavior would require to demonstrate the absence of relevant antecedent stimuli, making a firm case for operant behavior deserving a separate term proved elusive (Domjan, 2016).

One of the arguments in this debate can be summarized as an argument from incredulity:how could a behaving system evolve that produces behavior unrelated to its environment, how could such organisms successfully coordinate their actions with the world in which they live (e.g., (Domjan, 2016)? Today, we know from various observations both in the field and in the laboratory, that organisms without the capability to generate spontaneous actions would likely not have survived long. Individual behavioral variability has been found to be ecologically advantageous in game theoretical studies (McNamara et al., 2004; Glimcher, 2005, Glimcher 2003; Glimcher and Rustichini, 2004; Brembs, 1996), in pursuit-evasion contests such as predator/prey interactions (“Protean Strategy”)(Grobstein, 1994; Driver and Humphries, 1988; Shultz and Dunbar, 2006; Miller, 1997), in exploration/foraging (Belanger and Willis, 1996; Hills et al., 2013; Humphries and Sims, 2014; Shlesinger, 2009; Reynolds et al., 2016), in mobbing attack patterns by birds (Humphries and Driver, 1970) and in the variation of male songbirds’ songs (Neuringer, 2004). Clearly, invariable behavior will be exploited (Miller, 1997; Jablon ski and Strausfeld, 2000; Jablonski and Strausfeld, 2001; Catania, 2009, Catania 2010,Catania 2008; Mitra et al., 2009; Corcoran et al., 2009) and leaves the animal helpless in unpredictable situations (Brembs, 2009b, Brembs 2010; Heisenberg, 1994).

Two particular cases out of the multitude of examples for the evolutionary benefits of spontaneous behavior deserve special mention, both because the neurobiological mechanisms underlying the behavior are particularly well understood and because these behaviors have been used in operant conditioning experiments.

#### 1.1.1 Aplysia feeding

The first example is that of the feeding behavior of the marine snail *Aplysia*. These animals feed primarily on different species of seaweed, the texture of which can range from mushy to too tough to eat. The animals use their radula, a tongue-like organ, to grasp food and push it into the esophagus. The radula typically consists of two halves which can be either protracted or retracted while they are either open or closed. During initial feeding attempts, the animal protracts its radula in the open state, then closes the two radula halves and then retracts the radula, hopefully with a large chunk of edible seaweed enclosed in it (ingestion). However, if the food-item is too tough to push it further into the esophagus, the animal can also protract the radula while it is still closed around the food, pushing it out of the mouth. The animal then retracts the radula in its open state (rejection).

There are two aspects to the spontaneous components of this behavior.

The first one concerns the spontaneous variability in the movement dynamics of the first attempts to grasp the food and transport it towards the gut. *Aplysia* initially generates highly variable biting and swallowing movements, in an exploratory phase of feeding. During this time, the animal is exploring the state space of the motor system in the buccal ganglia controlling the movements of the radula. The animal is trying out which movements are most effective in term of transporting food into the esophagus (Lum et al., 2005; Horn et al., 2004). Reafferent (operant) feedback from dopaminergic fibers in the esophageal nerve instruct the buccal ganglia which movements were more or less effective (Nargeot et al., 1999c). This initial exploratory variability happens with all novel food items, it can thus be said to be independent of the nature of the food stimulus.

The second aspect concerns biting behavior in the absence of any food. Hungry *Aplysia* will spontaneously emit bites in the complete absence of any food stimulus. Clearly, it is always possible to argue that some undefined stimulus, commonly insufficient to elicit biting, becomes salient enough to trigger biting when the animal is hungry enough. However, in addition to any perceived implausibility of this argument, there is experimental evidence of neurons which have evolved precisely to generate spontaneous behaviors. Isolated buccal ganglia of hungry *Aplysia* will continue to generate the neural programs controlling the radula in the intact animal, when placed in a petri dish with suitable medium (Nargeot et al., 1999b, 1999a). These buccal motor programs (BMPs) occur spontaneously in the most explicit of terms:all sensory organs have been cut away, there cannot be any triggering stimuli in the petri dish. Moreover, the timing of BMPs is not rhythmic, the type of BMPs varies between ingestion -and rejection-like patterns and the intra-BMP dynamics vary from BMP to BMP. In other words, the *in vitro* experiments validate the *in vivo* work on spontaneous, exploratory biting behavior in *Aplysia.*

Physiological work on the neurons involved in generating BMPs identified a class of neurons that appears to have evolved to generate crucial aspects of this spontaneity (reviewed in (Nargeot and Simmers, 2012). Among the different classes of neurons controlling radula movements are two in particular which are relevant here. One of them, let’ s call them “ *What*” neurons, fires during the BMP and determine what type (ingestion-or rejection-like) of BMP will be produced, by controlling the timing of activity in the closure motor neuron. A second class of neurons, let’ s call them “ *When* neurons, fires right before a BMP can be observed in the motor nerves. Experimentally stimulating any of these *When* neurons will lead to the recording of a consecutive BMP in the motor neurons immediately following the stimulation. Thus, while the *What* neurons determine the type of behavior, the *When* neurons determine when a given behavior is going to be emitted. Part of the variability in the intra-BMP temporal dynamics comes from each BMP being initiated by a different neuron in the *When* class. One reason for the variability in which of the *When* neurons will start a BMP lies in the weak coupling between these neurons, preventing a stereotyped sequence of activity from emerging between the *When* neurons. A second crucial contribution to this spontaneous variability besides the weak coupling between them is the capability of individual *When* neurons to generate spontaneous bursts of activity even in the complete absence of input from other neurons. In contrast to the canonical neurons commonly studied, tonically stimulating an experimentally isolated *When* neuron to near its firing threshold leads to arrhythmic firing of the neuron (Nargeot and Simmers, 2012). The underlying molecular mechanisms are just being discovered and involve mathematically unstable, nonlinear calcium dynamics (Bedecarats et al., 2015). In the context of what roles these neurons play in the control of spontaneous behavior, it is straightforward to argue that these calcium dynamics have evolved to support ecologically relevant behavioral spontaneity.

In an analogue of the operant feedback described above, one can use any of these spontaneously generated BMPs and pair them with contingent stimulation of the esophageal nerve (Nargeot et al., 1999b, 1999a). This experiment thus establishes an *in vitro* operant conditioning paradigm out of a behavioral observation in freely behaving animals. This fortunate combination of intact and *in vitro* experiments, together with the exquisite physiological accessibility of *Aplysia*, allows for an unmatched rigor in the study of the learning processes actually taking place in the nervous system during operant conditioning. Perhaps not surprisingly, one such process directly affects the spontaneity of the behavior. More on that below.

Taken together, evidence both from intact animals and deafferentiated nervous systems demonstrates which aspects and components of *Aplysia* feeding behaviors are spontaneous in nature and what ultimate and proximate functions this spontaneity serves.

#### 1.1.2 *Drosophila* flight

*Drosophila* fruit flies with one injured wing are perfectly capable of flying straight, provided the injury is not too severe to prevent flight completely. This must seem like a remarkable feat to anyone steeped in the literature on how fixed optomotor reflexes control straight flight in *Drosophila*. Unless the animal is born with the knowledge of exactly which kind and amount of wing damage leads to which effect on torque kinematics, there must be some reafferent feedback that instructs the flies on the effectiveness of their turning maneuvers. Several observations support the conclusion that flying straight is primarily an operant behavior (reviewed in, e.g., Brembs, 2009b). For instance, one can experimentally eliminate all optomotor responses (without rendering them blind) and the flies are still able to fly straight. Crucially, these manipulated flies can do this even if the feedback between their behavior and the environment is experimentally reversed, i.e., left turning attempts lead to visual feedback that suggests a right turn and *vice versa* (similar to inversion goggles in humans). These experiments are done with the flies tethered to a torque meter, which measures the yaw torque of the flies without them actually rotating in space. Surrounded by a visual panorama that can be rotated around the flies instead, the experimenter has exquisite control over the flies’ stimulus situation. Tethered optomotor-disabled flies manage to keep the visual panorama from rotating (i.e., fly straight) no matter how the rotation is coupled to their torque behavior. This is in striking difference to unmanipulated flies which need a very long time until they manage to keep the panorama steady with inverted feedback (but do so within milliseconds in regular coupling). Apparently, the inborn optomotor responses impede the wild type flies in this experiment, while the symmetrical behavior of the optomotor-impaired flies is evidence that they must use operant behavior to minimize the amount of arena rotation in the absence of any sensory system telling them the direction of rotation (Wolf and Heisenberg, 1986). As with the example of *Aplysia* above, with these flies being partially deafferentiated, any of their turning attempts must be spontaneous as there are no optomotor stimuli being perceived that could elicit a turning response.

Another experiment brings us back to the ability of the flies to compensate for one-sided wing injury. The kind of turning maneuvers the optomotor-impaired flies are using to control their visual feedback can also be observed in tethered wild type animals in a completely uniform environment. Similarly to the manipulated flies, there are no visual stimuli known to elicit turning attempts. Great care is taken to ensure that all the stimuli that are present, of any modality, are as constant as experimentally possible. In fact, from the experiments with external stimuli present, it is known that the remaining, constant stimuli do not exert any detectable effect on top of the explicit stimuli. Again, without complete experimental deafferentiation, it is impossible to be sure that these sensory deprived animals are not responding to otherwise unphysiological stimuli that bear no relation to flight under normal circumstances. One would assume that these stimuli would be occurring randomly, without any specific structure or pattern. In that case, the temporal dynamics of the attempted turning behavior of such deprived flies should be reminiscent of random noise. However, the temporal dynamics of these flies is much more reminiscent of the mathematically unstable, nonlinear process discovered in *Aplysia.* In a recent transgenic screen, we are beginning to identify candidate circuits comprising the neurons that may be responsible for this nonlinear signature. Interestingly, these structures appear to be located in the same brain regions as those associated with the temporal structure of walking behavior (Martin et al., 2001). Recent optophysiological work on zebrafish is also beginning to identify the brain regions in the vertebrate brain where the temporal structure spontaneous turning movements is controlled (Dunn et al., 2016).

Instead of visual feedback, one can use heat as a feedback in the tethered flight experiment and let the fly use its turning attempts to control the punishing heat beam (in a completely homogeneuous visual environment). When, e.g., left turning attempts are punished, the fly shifts its baseline torque towards the unpunished (e.g., right) direction, without eliminating superimposed bidirectional fluctuations in torque, even after the heat is permanently switched off (Wolf and Heisenberg, 1991). This capability is precisely what would be required to compensate the reduced torque of a damaged wing:a shift in the baseline torque output, operantly matched to the (visual) feedback, without eliminating the possibility for left and right turns from this new ‘straight-flight’ baseline.

Taken together, converging evidence from *Aplysia* and *Drosophila* suggests that spontaneous behavior is controlled by an evolutionary conserved mechanism that relies on unstable nonlinearities to explore the state space of the motor system in search for favorable behavioral outputs.

## 2 Behavior is likely never a “response”

Given the spontaneous components in even highly stereotyped behaviors such as feeding or optomotor control, one is tempted to examine what one may consider the simplest stimulus-response systems for evidence of spontaneity.

### 2.1 Phototaxis in insects

One could consider the movement of insects towards a light source (phototaxis) as one such system. The fact that insects get trapped at windows or the proverbial moth that dies in a candle flame are iconoclastic examples of a rigid stimulus-response organization of this behavior. But does insect phototaxis stand up to scientific scrutiny? Already a hundred years ago, observations by McEwen (1918) suggested that this behavior may be more complex than one would at first expect. McEwen found that only startled flies would walk towards the light in his small tube. Sitting flies did not seem to find a light very attractive, suggesting that more than just light hitting the retina must be responsible for triggering phototaxis. He also observed that flies with clipped wings do not approach the light anymore, even when compared to walking, intact flies. This observation was later confirmed in a different experiment, where the wings were not only clipped but also genetically rendered useless for flight (Benzer, 1967). Further experimentation revealed that at least for insects, the term ‘phototaxis’ may be inappropriate (Gorostiza et al., 2015). Rather than just affecting the approach of a light source, the flies’ ability to fly affected their light/dark preference across several different behavioral tests, none of which tested phototaxis, but forced the flies to choose between more or less bright stimuli. If flying ability was compromised only temporarily, the flies’ photopreference reversed concomitantly. Neuronal activity in circuits expressing dopamine and octopamine, respectively, doubly dissociated in this case of behavioral flexibility (Gorostiza et al., 2015):activity in octopaminergic neurons was necessary and sufficient to shift the flies’ preference towards darkness, while activity in dopaminergic neurons was necessary and sufficient to shift the preference towards brightness. The involvement of these biogenic amines suggests that valuation of stimuli may play a role in the flies’ shifts in photopreference. Apparently, flies monitor their ability to fly, and the outcome of this evaluation exerts a fundamental effect on action selection - including, but not exclusively in phototaxis experiments. This work suggests that even innate preferences which appear simple and hard-wired, such as those expressed in classic phototaxis experiments, comprise a value-driven decision-making stage, negotiating external stimuli with the animal’s internal state and likely other factors as well, before an action is selected. This endows the animal with the possibility to decide, for example, when it is better to move towards the light or hide in the shadows. Moreover, the fact that flies adapt their photopreference in accordance with their flying ability shows that flies have the cognitive tools required to evaluate the capability to perform an action and to let that evaluation impact other actions-an observation reminiscent of meta-cognition.

Thus, what appears to be a simple response to light, actually contains at least one decision-making stage which negotiates and weighs several different factors before selecting an action. One may even argue that what was commonly described as a taxis, in this case at least, only appears as a simple taxis at a superficial glance. Once one peers into the neurobiology of the behavior, the usage of terms like ‘response’ or ‘taxis’ appears inadequate.

### 2.2 Knee jerk reflexes in mammals

A similarly iconoclastic input-output system is the “knee jerk” class of spinal reflexes. One can hardly imagine a simpler system:the 1a afferents send the excitation from the stretched muscle spindles to the ϒ motor neurons which activate the muscle that flexes the leg. On the surface, this seems to be an even simpler and more obviously feed-forward case than phototaxis in insects:two neurons, one synapse, input from the sensory neuron leads to output from the motor neuron and behavior. However, even there, upon closer examination, the seemingly dominant stimulus-response organization starts to collapse.

Implanting cuff electrodes on the mammalian posterior tibial nerve as well as electrodes recording the electromyograms from the soleus muscle allows for longterm recordings of the electrical analog of the stretch reflex, the H-reflex (or Hoffmann’ s reflex). The simple textbook case of the knee jerk reflex implies all-or-nothing responses to identical stretch stimuli. However, triggering this reflex over hours, days or weeks reveals multiple timescales of variability in the amplitude of the H-reflex. Moreover, making a food or water reward contingent on larger (or smaller, respectively) reflex amplitudes than baselines averages leads to an increase (or decrease, respectively) of the reflex amplitude over the course of a few days (Wolpaw, 2010; Chen and Wolpaw, 1996; Wolpaw and Chen, 2006; Carp et al., 2006; Thompson and Wolpaw, 2014a, Thompson and Wolpaw, 2014b). One could say that, conceptually, this procedure is the opposite of an omission schedule:to eliminate potential operant components in classical conditioning experiments, an omission schedule leaves out the unconditioned stimulus (often food or water), whenever the conditioned reflex was produced. In operant conditioning of the H-reflex, only those reflexes are rewarded that reach the target amplitude. In this case, always the same electrical stimulus is eliciting the reflex, so the behavioral variability must be either due to internal processes in the animal or due to unrelated stimuli in the environment. It is difficult to imagine that, for instance, a certain corner of the experimental chamber consistently increases reflex-amplitudes throughout the nervous system, such that the animal would always seek this corner to increase its amplitudes during conditioning. On the contrary, such operant conditioning in humans is used to improve rehabilitation after spinal cord injury (Thompson and Wolpaw, 2015, Thompson and Wolpaw, 2014a), such that lasting changes in reflex amplitude must be brought about by processes that are independent from the current environment of the patients. In fact, the function of the spontaneous variability in reflex amplitude is quite well understood:the spinal reflexes constantly adapt to the environment in which vertebrates walk. They accomplish this feat by constantly changing their amplitude and evaluating the sensory consequences of these actions. In other words, even spinal reflexes are using spontaneous actions to constantly explore the state space of the motor system in order to find the most suitable behavioral output. In this process, the spinal reflexes rely on networks that span the cortico-spinal tract all the way into the cerebellum and motor cortex. Without these reflexes using these networks to constantly probing the environment’s responses to their spontaneous actions, we would not be able to walk up the stairs at the end of the hallway or climb down a mountain after we reached the summit.

Thus, the textbook knee jerk reflex only exists in its textbook form if the slice of the spinal cord that contains its synapses is removed from the rest of the nervous system. However, this isolated state prevents understanding of the function of spinal reflexes for locomotion. Describing these spinal reflexes as rigidly responding to stretch stimuli seems at best inadequate and at worst misleading, considering the large-scale networks within which this sensory-motor synapse is embedded and the complex, operant processes these circuits in fact mediate.

### 2.3 Phototaxis in a polychaete larva

A less iconoclastic, but perhaps yet simpler and likely one of the most archaic of these stimulus-response systems, is also considered a model for the last common ancestor of vertebrates and invertebrates, the ‘Urbilaterian’:the planktonic larvae of a marine polychaete worm, *Platynereis dumerilii.* These eggshaped creatures use a band of ciliated cells around their body to locomote, in the first half of larval development preferentially towards the surface. One factor in this upward movement is positive phototaxis. When a light is switched on at one end of a chamber filled with *P. dumerilli* larvae, they all start swimming towards it (Je kely et al., 2008). That these animals are capable of phototaxis is remarkable as their nervous system does not feature any interneurons. Their light-sensitive neurons make direct synaptic connections with the ciliated cells that propel the animal. Importantly, the ciliated cells are constantly active, propelling the animals sometimes in this directions, sometimes in that. When light hits one of their two ‘eyes’, the synaptic connection between the light-sensitive neuron and the ciliated cells inhibits the ciliated cells on the side where the light was perceived (Je kely et al., 2008). Like a rower who stops rowing on one side, the animals then rotate towards the side where the light came from. If they keep rotating, light will hit their other eye, leading, again, to a rotation towards the light. This physiological understanding of the biological mechanisms underlying phototaxis in *P. dumerilii* larvae is necessary for the insight that what appears as an external stimulus *eliciting* an until then inactive behavior is organized rather in the reverse fashion:these animals are constantly exploring their environment in the search for light, constantly changing directions. The external stimulus is then perceived as feedback from this exploratory, spontaneous behavior. In stark contrast of the implied activation of a response, the stimulus then only serves to *eliminate* a portion of the ongoing behavioral repertoire.

Thus, it appears as if behavior in a model for the Urbilaterian is organized in a fashion antithetical to the stimulus-response organization often assumed for nervous systems in general. These animals are first generating ongoing, random(-like), exploratory behavior that is modulated by subsequent reafferent feedback. This discovery may constitute the simplest, maybe even the earliest instantiation of an operant organization of behavior:generating a spontaneous action first and then evaluating its outcomes. If that were the case, ‘responses’, if they actually exist, may only be rare and highly specialized, evolutionarily relatively late adaptations. In this view, the general concept of a stimulus-response organization of behavior is largely due to a combination of selection bias in which animal models and experiments are chosen for study and an inevitably superficial observation of the behavior.

### 2.4 Olfactory Reversal in Nematodes

If a stimulus-response organization of behavior were a particular evolutionary adaptation, e.g., evolved to speed up action selection in predictable situations, perhaps one can find examples of them in an animal model where we have an indication that evolution may have streamlined the pathways from stimuli to responses. One of the most well-studied genetic model organisms and so far the only adult animal with a complete connectome of its nervous system is the nematode worm *Caenorhabditis elegans* with its 302 neurons. The *C. elegans* connectome is dominated by feed-forward connections from sensory neurons to motor neurons (Qian et al., 2011), so maybe the nervous system of this nematode is a promising candidate to find ‘ responses’ in the literal meaning of the word.

One well-characterized behavior in this nematode is reversal behavior. It occurs whenever the animal encounters aversive stimuli, such as certain odors. The circuit controlling this behavior can be described with just four neurons, their 44 chemical connections and their electrical synapses. A central component of the system is a neuron called AVA. When AVA is active, the animal reverses its course. Sensory input to this neuron is provided by an olfactory neuron, AWC. For instance, if AWC is stimulated by an attractive odorant, it stops firing, such that AVA loses excitatory input and also stops firing, making reversals less likely. Conversely, activating AWC either experimentally or with an aversive odor increases the probability of reversals by synaptically activating AVA (Gordus et al., 2015). Two additional neurons are involved in this circuit, AIB and RIM, and the characterization of their role in the circuit is crucial for understanding the organization of olfactory mediated reversal behavior in *C. elegans*.

The first interesting observation from the circuit connectivity is that there are more connections from the sensory AWC neuron to the AIB interneuron than to the reversal neuron AVA. This is unexpected, if the main function of nervous systems were to relay sensory information to motor centers. Imaging this circuit in immobilized worms in the absence of any stimuli, reveals a complex patterns of correlated activity in all neurons. Interestingly, the neurons exhibit a sort of binary activity state, that for the most part is either on (neuron is active) or off (neuron is inactive). Quantifying the activity fluctuations in this circuit, one finds that there are three main states (of the eight theoretically possible) the circuit is commonly found in:just over 60% of the time the system is in ‘all on’, roughly 20% is ‘all off’ and for the remaining 20% it is in the state ‘only AIB on’. This observation yields two insights:For one, even without any stimulation at all, these network dynamics can generate spontaneous reversals without requiring any sensory input. Second, each olfactory stimulus reaching AWC will interact with the state the circuit currently happens to be in, rather than arriving in a quiescent circuit and triggering some neural activity that was not there before. The behavioral consequence of this interaction is not only the occurrence of spontaneous reversals, but also the occurrence of ‘spontaneous’ non-reversals in the presence of an aversive odor. In other words, the reversal circuit is probabilistic and without observing the nervous system of the worm, it is impossible to tell how spontaneous the observed behavior actually is.

Experimentally silencing either one or both of the interneurons in this circuit reveals that the role of AIB and RIM is to increase the variability of the reversal circuit. While the input to the circuit from the olfactory neuron AWC is always very precise and predictable if, e.g., an odor is presented, the activity of the reversal circuit always varies significantly and this variability is reduced if AIB or RIM (or both) are silenced (Gordus et al., 2015). This discovery makes an excellent case for RIM and AIB being incorporated into the reversal circuit specifically to inject much needed variability into an otherwise maladaptively deterministic reversal circuit. Surprisingly, even though the feed-forward connections dominate the connectivity also in this little circuit, the variability provided by the feed-back connections dominate an adaptive feature of the behavior, its variability. This work adds *C. elegans* to the elongating list of animals, whose nervous systems are organized such that ongoing activity is merely modulated by external stimuli. In the nematode case, it appears that out of the four neurons comprised in this circuit, two exist for the sole reason to mitigate the effects that stimuli have on the behavior of the animal, in order to make the animal more autonomous with regard to its environment. If an animal with only 302 neurons, which, as in all other animals, make up the most energetically costly tissue, devotes 50% of a circuit to counter the effects of stimulus-response connections in its nervous system, then the implications of this discovery for the organization of behavior in animals generally cannot be underestimated.

Thus, contrary to the idea that a connectome dominated by feed-forward connections from sensory to motor areas implies that it mainly computes motor output from sensory input, also the nervous system of *C. elegans* is best characterized by constantly changing, ongoing activity, much like many other nervous systems previously studied in this regard. It seems that even a numerically small feed-back component provides a fundamental contribution to the overall architecture even of such feed-forward-dominated networks. What does this mean for brains, such as those of mammals, whose neuroanatomy appears to be dominated by feed-back loops?

### 2.5 Escape responses

Few behaviors are as obviously under evolutionary selection pressures as escape responses:the anti-predator behavior of prey in the presence of predators. Perhaps not surprisingly, this class of highly refined behaviors is particularly well-studied because of their reproducibility in the lab. Be it the C-start response in fish, mediated by the largest mammalian neuron, the Mauthner cell (Korn and Faber, 2005; Schuster, 2012), the squid escape response mediated by their giant fiber system (Young, 1938), the crayfish giant fiber system that allows the animal to quickly propel itself out of harm’ s way by flipping its tail (Herberholz and Marquart, 2012) or the fly escape circuit also mediated by giant fibers originating in the optic lobes and making direct connections with the motor neurons that lead to both the raising of the wings and to the jump response of the legs (Hammond and O’ Shea, 2007a; Card and Dickinson, 2008b, Card and Dickinson 2008a). All of these behaviors are easily observed in the lab, are mediated by conspicuous and easy to manipulate giant neurons and have, over the decades, shaped the way neuroscientists all over the world once conceptualized how behavior is organized:eliciting stimulus in, followed by behavioral response out (e.g., “brain function is ultimately best understood in terms of input/output transformations” (Mauk, 2000). However, studies in behavioral ecology and ethology have revealed that these famous behaviors are liable to exploitation (see, e.g., (Catania, 2009,Catania 2010; Jablon ski and Strausfeld, 2000; Jablonski and Strausfeld, 2001) and/or are often more complex and flexible than initially thought (Card and Dickinson, 2008a, 2008b; 2007a,Hammond and O’ Shea 2007b; Herberholz and Marquart, 2012; Fotowat and Gabbiani, 2007). Searching for more escape behaviors under natural conditions in order to compare them with the better known laboratory examples, it was discovered that many escape behaviors contain elements of variability in order to make the escape trajectory less predictable for the predator (e.g., (Royan et al., 2010; Domenici et al., 2008; Bateman and Fleming, 2014; Driver and Humphries, 1988; Humphries and Driver, 1970; Guerin and Neil, 2015; Highcock and Carter, 2014). This work demonstrates the adaptive value of such “protean” escape strategies and suggests that unpredictable prey is not only more difficult to catch, but is also capable of injecting additional unpredictability into their behavior in the presence of predators. It thus appears as if the technically advantageous property of being highly reproducible not only renders an escape behavior liable to exploitation in the wild, but has also introduced a bias in neuroscience and psychology:reproducible laboratory behaviors are rarely representative of behaviors more generally, which tend to be much more variable and contain a larger degree of unpredictable spontaneity.

In the light of such data, it is tempting to postulate that the stimulus-response concept of animal behavior is little more than a laboratory artefact introduced into those sections of psychology and neuroscience that have isolated themselves from non-laboratory behavior. Instead, converging data from multiple fields, model and non-model organisms in the laboratory and the wild suggests that at least all bilaterians have evolved the capability to inject a controlled amount of variability into their behavior in order to accomplish adaptive behavioral choice under a variety of situations. The influence of this “random number generator” ranges from changing minute aspects in the temporal dynamics of very simple behaviors to drawing from a set of different behaviors. The hallmark of such behavioral variability, the feature that confers its adaptive value is its independence from the environment, i.e., its spontaneity. For instance, the presence of a predator may increase a prey’ s behavioral variability, but not a specific escape trajectory or direction. In a novel environment, an animal may increase the variability of its foraging or exploratory behavior, but the environment cannot be used to predict the direction or duration of the foraging bout or the timing of the next exploratory behavior (see, e.g. (Jacobs and Menzel, 2014) for a similar argument in a different research question).

## 3 The interaction between ‘self’ and ‘non-self’

The observations discussed so far have led to a much more refined and sophisticated picture of how nervous systems organize behavior than can be circumscribed by the ‘emitted’ and ‘elicited’ dichotomy. From a neurobiological perspective, stimuli interact with and modulate ongoing activity in the nervous system, rather than triggering always the same cascades of neural activity in a more or less quiescent brain. This insight entails a number of important consequences.

For one, as behavior is based on neural activity, it is ongoing and continuous. The impression that behavior can be chunked into discrete acts (i.e., “units of behavior”) are possibly due to both our tendency to categorize even continuous phenomena as well as the scale-free nature of neural activity, allowing us to find temporal patterns in behavior at different time scales.

Another consequence is that sensory information only needs to instruct the animal as to which kind of behaviors to favor over others. Which specific movements or actions are to be generated do not always have to be explicitly specified. Exploratory ‘ trying out’ with efficient feedback evaluation (i.e., operant behavior) is sufficient to quickly and reliably find the behavior that is best suited to solve the task at hand.

However, it is clear that some stimuli exert a more specific effect on the ongoing activity in the brain than others. There appears to be a continuum in the urgency or salience of events in an animal’ s environment, at one extreme end of which are stimuli so potent, that they appear to trigger a behavioral response, at least to the casual or superficial observer. Obviously, the coupling between a stimulus and an organisms behavior can vary from very loose to very tight both in terms of temporal coupling as in terms of which stimuli are followed by which behaviors, and *vice versa*. Most often, speed and efficiency are correlated with a tight environmental coupling, not only for stereotypic behaviors such as, e.g. escape responses:the well-trained squirrel will much more quickly crack the nut than the naV ve one. However, fast, efficient and ultimately stereotypic behaviors are inflexible and liable to exploitation. Inasmuch as operant conditioning leads to the selection of efficient behavior and later to stereotypization of the behavior in a process called habit formation, operant processes are central in the animal’ s quest to trade off efficiency for flexibility and unpredictability. This trade-off is essential for the survival and procreation of every organism and likely one of the most important ultimate causations behind adaptive behavioral choice.

Of course, there are ongoing processes inside the organism that can be ranked on a similar scale from hardly noticeable to directly influencing behavior. Hunger or circadian rhythms have at least as potent an effect on behavior as the presence of food, a predator or a potential mating partner. It is the complex interaction of processes generated by the animal itself (among which spontaneous behavior can be found) with processes generated by events outside of the animal that ultimately manifests itself as observable behavior. Importantly, the fact that we cannot tickle ourselves suggests that this distinction between self and non-self may be much more fundamental than one would at first assume. In their “ reafference principle” von Holst and Mittelsta dt famously proposed a mechanism by which animals use efference copies to distinguish between stimuli that are under their control (reafference) from those that are not (exafference)(von Holst and Mittelstaedt, 1950). The reafference principle is a fundamentally operant process by which animals not only distinguish between the stimuli they can control and those they cannot:it is also the beginning of the operant process by which animals learn to bring novel stimuli under their behavioral control.

These considerations bring us back to another fundamental question Skinner formulated in the 30s of the last century. Given the multitude of learning processes engaged during operant conditioning experiments, how many of them are similar to those taking place in classical conditioning (i.e., stimulus substitution) and how many, if any, are fundamentally different. Skinner himself lamented that he could not remove the lever his rats were pressing to eliminate one of the confounding factors (Skinner, 1935). Modern behavioral neurobiology in genetic model organisms have designed experiments specifically to address this problem and have discovered that the distinction between self and non-self is central to a purely operant learning mechanism.

## 4 Mechanisms of Plasticity in Operant Conditioning

### 4.1 Isolating the components:self-learning

As Skinner noted, rats need a lever to press and thus they may learn about the food-predicting properties of the lever. Therefore, this experiment is not ideal for studying the neurobiology underlying operant learning processes. Any memory trace found in the brain cannot be unambiguously attributed to the mechanism engaged when learning about the lever or to the learning about the behavior required to press the lever. Therefore, preparations had to be developed without such environmental ‘ contamination’.One such preparation is tethered *Drosophila* at the torque meter as described above (Brembs, 2009b; Heisenberg et al., 2001; Wolf and Heisenberg, 1991; Heisenberg, 1994; Wolf et al., 1992; Heisenberg and Wolf, 1984; Wolf and Heisenberg, 1986). In the setup where the flies learn to control a punishing heat beam with their yaw torque in a homogeneous environment with no directional cues, the only contingency present is that between the behavior and the reinforce/punisher. In this experiment, it is thus possible to isolate the operant contingency from all environmental contaminations. One may term such experiments ‘ pure’ operant conditioning experiments.

Using mutant, wildtype and transgenic animals, it was discovered that the canonical, cAMP-dependent synaptic plasticity pathway known from other learning experiments was not involved in this type of learning, but manipulating protein kinase C (PKC) signaling abolished learning in this paradigm completely (Brembs and Plendl, 2008). Apparently, the common, evolutionary conserved mechanisms, discovered in classical conditioning experiments are not required for this pure operant learning. Instead, the animal has to rely on a different biochemical process if it is asked to learn about its own behavior, in the absence of any external cues. With regard to the fundamental distinction between self and non-self discussed above, the content of the learning process is fundamentally about the animal’ s own behavior. Thus, one may call this process self-learning, i.e., the process by which value is assigned to a specific action or movement, such as the heat is assigning positive or negative value to left or right turning, respectively, in this experiment (Colomb and Brembs, 2010).

In the search for other components of the biological processes underlying self-learning, one may again refer to Skinner. This time his claim that language acquisition constituted a form of operant learning (Skinner, 1957):first exploratory, highly variable actions are being initiated (i.e., babbling) and then sensory feedback shapes the initiation of future behavior, reducing its variability (i.e., language). In what may appear at first to be a very superficial analogy, “ mere homonyms, with at most a vague similarity of meaning” (Chomsky, 1959), operant self-learning in tethered flying *Drosophila* mimics these features of vocal learning:the animal first initiates highly variable, exploratory actions, then sensory feedback shapes the initiation of future behavior, reducing its variability. The Forkhead Box P2 (FOXP2) transcription factor is the first gene discovered to be involved in the development of speech and language (Fisher and Scharff, 2009; Lai et al., 2001). Importantly, the avian orthologue is also involved in song learning in birds (Scharff and Haesler, 2005; Haesler et al., 2007; Schulz et al., 2010), which has also been described as an operant behavior (Marler, 1991). The four different FoxP genes in vertebrates probably arose from serial duplications of a single ancestral FoxP gene after the separation from the invertebrate clades. The invertebrate FoxP orthologue corresponds most closely to the ancestral form of the gene at the base of the bilateria (Santos et al., 2011), thus lending itself to investigating the depth of the functional conservation among the members of the FoxP gene family. Mutant analysis and RNAi-mediated knockdown of the *Drosophila* orthologue, *dFoxP*, revealed its necessity specifically for self-learning (Mendoza et al., 2014), a phenocopy of the PKC manipulations described above. Thus, an excellent candidate for the ancestral function of FoxP genes, several of which are involved in acquiring the speech component of language in humans, is specifically involved in the isolated self-learning component of operant conditioning, but not in other forms of learning in *Drosophila*. These results are consistent with the hypothesis that the FoxP-dependent component of language evolved from an ancestral operant self-learning mechanism. The homology between invertebrate FoxP and its descendant genes, together with the similarities between habit formation and birdsong crystallization (Costa, 2011) (see also the Chapter on bird learning) prompts the postulation that habit formation in vertebrates may also be engaging this same mechanism.

Parallel developments to isolate the operant component have been made in the sea slug *Aplysia* (Nargeot et al., 1997; Nargeot, 2002; Nargeot et al., 2007, 2009; Nargeot and Simmers, 2010; Brembs et al., 2002; Lorenzetti et al., 2008, 2006; Nargeot et al., 1999c, 1999b, 1999a). As described above, reward signals were made contingent on spontaneous biting behavior, either in the intact animal or in isolated buccal ganglia. This procedure also excluded any external stimuli from contaminating this pure operant experiment, leading to self-learning. Such conditioning brings about two changes in the behavior of the animals/ganglia:both the total number of bites/BMPs is increased and the frequency of the rewarded behavior increases, relative to the other feeding behaviors. Electrophysiological studies discovered learning-related changes both in *When* and in *What* neurons. While the *What* neurons changed their excitability such that the behavior they promoted became more frequent (Brembs et al., 2002; Nargeot et al., 1999b,1999a), the *When* neurons increased their electrical couplings as well as their excitability (Nargeot et al., 2009), such that not only the total number of behaviors increased, but they also became more stereotyped, losing the variability described above for the naV ve animals in which the coupling between the *When* neurons was comparatively weak (Nargeot and Simmers, 2012). The self-learning mechanism in *Aplysia* hence entails direct modifications of the very neurons controlling the type of behavior being generated as well as the variability of the behavior. In this way, we are starting to unravel the neuronal mechanisms behind selection by consequences (Skinner, 1981) (see also the chapter by Aaron Blaisdell). These mechanisms appear to be evolutionary conserved as well, raising the possibility of a common, specifically operant learning mechanism for all bilaterians. Also in *Aplysia*, as in *Drosophila*, the canonical cAMP-dependent learning pathway discovered in classical conditioning is not involved, but PKC manipulations impair self-learning (Lorenzetti et al., 2008). Given that PKC is also involved in song-learning in birds (Yoshida et al., 2003; Sakaguchi and Yamaguchi, 1997), as is FoxP2, there is now strong evidence for such a conserved self-learning mechanism.

These results entail that there exists a dedicated biological mechanism that only occurs in neurons that are involved in actions and not in those processing the environment. With regard to the early questions of Skinner and his contemporaries, we can now say that we have learned that the different associative processes mediating the content of learning (“what is learned”) in operant conditioning are mediated by different biological mechanisms:the molecular machinery involved in operant self-learning (i.e., PKC and FoxP to date) does not appear to be involved in any of the other types of learning studied so far. This insight was made possible by separating and isolating the self-learning process from any other processes that may take place during operant conditioning. Adding some of these components back to the experiment reveals interactions between these components that have a direct relation to the central efficiency/flexibility trade-off underlying adaptive behavioral choice discussed above.

### 4.2 Combining the components:composite learning

Once the operant component has been isolated in ‘pure’ operant learning experiments in which only self-learning can occur, it is comparatively easy to add a predictive stimulus and compare the resulting ‘composite’ situation with the ‘pure’ experiment. For instance, in tethered *Drosophila,* whenever the direction of turning maneuvers changes (e.g., from left to right turning attempts), the entire visual field of the fly instantaneously turns from one color (say, green) to another (e.g., blue). Because now the colors change both with the yaw torque and the heat, the fly has the additional option to learn that one of the colors signals heat, and not only that its own behavior contriols the heat. This situation is analogous to how rats in Skinner-boxes may learn that the depressed lever signals food (and the undepressed lever no food) as well as that their lever-pressing is predictive of the food reward. In contrast to the ‘pure’ experiment, this situation now requires the canonical cAMP cascade discovered in classical conditioning and is independent of any PKC signaling (Brembs and Plendl, 2008) or *dFoxP* function (Mendoza et al., 2014). Apparently, even though the experiment is just as operant as before the colors were added (in fact, if the flies were able to close their eyes to the colors, it would be the exact same, ‘pure’ experiment), now seemingly ‘classical’ learning mechanisms are engaged. This result suggests that as soon as learning of external stimuli becomes possible, these will be learned preferentially over behavioral cues, even in otherwise completely operant learning situations. As the content of these learning processes is fundamentally non-self, one may call these processes ‘world-learning’, i.e., the process by which value is assigned to external stimuli, such as, for instance, ‘green’ being assigned an aversive value and blue an appetitive value in this experiment (Colomb and Brembs, 2010). In contrast to self-learning, world-learning can occur in both operant and classical learning experiments, depending on the operant control of the predictive stimuli (this will become a separate point of discussion below). Thus, the difference between operant and classical conditioning lies not in the procedural differences between the two experiments, it lies in the content of the memory being formed:its content (i.e., ‘ what is learned’) decides which learning processes are engaged in the nervous system. How this content is learned, appears to be less relevant.

To ask whether the colors have been learned independently from the behavior with which they were learned, one can test the flies’ preference of the unpunished color with an orthogonal behavior to that used during training. After all, to solve the ‘ composite’ situation, it is sufficient for the flies to learn that one of the colors is associated with the heat and then use whatever behavior necessary to avoid this color. Indeed, flies can avoid the punished color even with an orthogonal behavior. Conversely, when tested for a preference in turning direction (i.e., without colors) after composite conditioning, the flies do not reveal any preference (Brembs, 2009a). In the most Pavlovian sense, the flies seem to learn the color-heat contingency independently of the behavior with which it was acquired. They only learn about the world around them, without leaving much of an indication that the behavioral decision-making circuitry itself has been significantly altered, even though, of course, the entire situation is still just as operant as without the stimuli (Brembs, 2009a; Brembs and Heisenberg, 2000). Apparently, there is an inhibitory interaction between the world-and the self-learning processes during operant composite conditioning, such that only the effects of the world-learning process can be detected after operant composite conditioning. For this inhibition to occur, it is not sufficient that the colors are merely present:flickering the colors unrelated to the heat does not inhibit self-learning and mutants which cannot learn the colors (but are not impaired in self-learning) do not show any sign of self-learning inhibition, even if the colors are predictive of the heat (unpublished observation). Thus, world-learning must be actively engaged, in order to inhibit self-learning.

To ask whether this inhibition is absolute or depends on other factors, such as, e.g. time, one can extend the training to twice the duration used in traditional operant conditioning experiments in *Drosophila.* Testing for world-and self-learning effects after such extended training reveals that the inhibition of self-learning by world-learning is time-dependent:after extended training, the flies prefer generating the previously unpunished turning direction, even in the absence of the colors. In what appears to be an analogy to vertebrate ‘ habit interference’, the flies can no longer express the preference for the previously punished color with an orthogonal behavior (Brembs, 2009a). Interestingly, *FoxP* mutant flies are impaired in this type of habit formation, indicating that the self-learning that is taking place in pure operant conditioning is indeed the very same self-learning process that is inhibited in composite conditioning. One may interpret these results such that habit formation requires repetition because it is inhibited by world-learning. After prolonged training, this inhibition is overcome and self-learning kicks in to form habits. In flies, a prominent neuropil, which is dispensable for both world and self-learning, is involved in the inhibition of self-learning:the mushroom-bodies (Brembs, 2009a).

Similar relationships have been observed in experiments with other animals. For instance, in navigation studies, relatively short training preferentially engages an allocentric strategy (the animal orients primarily according to environmental cues), while longer training induced an egocentric strategy (the animals performed the same sequence of movements)(Packard and McGaugh, 1996; Hicks, 1964; Ritchie et al., 1950; Tolman et al., 1947, 1946). The analogy to world-and self-learning is striking. The terminology of world-and self-learning itself was inspired by analogous developments in another research field (Berniker and Kording, 2008). There is a third field in which analogous results have been obtained. In experiments with rodents in operant chambers, extended training abolishes sensitivity to reinforcer devaluation by the process of habit formation which transforms goal-directed actions to habitual responses (Yin and Knowlton, 2006).

Besides the inhibitory interaction between world-and self-learning, there is a second interaction that can only be observed in composite conditioning. These experiments take advantage of the fact the world-learning can occur in operant conditioning experiments (i.e., if they are composite experiments) and always occurs in classical conditioning experiments. If fly learning in composite situations is compared to situations where the same stimulus was trained classically, it is routinely observed that the stimuli are learned faster and to a higher level in the composite, than in the classical situation, even if the sensory input during training was identical between groups (Brembs and Heisenberg, 2000; Brembs and Wiener, 2006). This observation is reminiscent of the generation-effect (“learning-by-doing”), i.e., the facilitation of world-learning by being in control of the stimuli which are to be learned (Thorndike, 1898; Slamecka and Graf, 1978; Kornell and Terrace, 2007; James, 1890; Baden-Powell, 1908). The mechanism by which this facilitation of world-learning occurs in composite conditioning compared to otherwise identical classical conditioning remain elusive. The only results so far are negative:none of the mutants and transgenes tested in the last two decades shows any deficit in the ‘generation effect’.

It is straightforward to hypothesize that the difference in world-learning rate between classical and operant composite conditioning is due to the stimuli being presented exafferently in classical conditioning and reafferently in operant situations. Efference copies are generally proposed as one mechanism animals use to distinguish between exafferent and reafferent stimuli. If efference copies are also used here to accomplish the generation effect in composite conditioning, it would entail that efference copies not necessarily always reduce the salience of reafferent stimuli as originally proposed (von Holst and Mittelstaedt, 1950), but can also enhance it, under certain circumstances, to accomplish a facilitation of world-learning. From these considerations it is a small step to postulate that when the relationship between a reafferent stimulus and the animal itself is concerned (such as in re-afferent self-motion signals, or in self-tickling attempts) then efference copies serve to reduce the salience of the stimuli. However, when the relationship among different reafferent stimuli is concerned, then efference copies serve to increase their salience compared to the same stimuli presented exafferently.

## 5 Conclusions

In contrast to the long-held assumption that the organization of behavior is best characterized as the perception of a sensory stimulus followed by appropriate response (i.e., “sensorimotor hypothesis”), recent converging evidence from multiple systems and fields of study instead suggests that both ancestral and extant general brain function is best described in operant terms. Rather than specifying precise behaviors, sensory information – if at all present – interacts with ongoing neural activity to instruct the organism which type of spontaneous, exploratory behavior to generate. Evaluating the ensuing reafferent feedback modifies the nervous system such that ongoing neural activity patterns become biased towards activity that has generated increased appetitive and decreased aversive feedback in the past. The neurobiological mechanisms underlying both the exploratory, spontaneous behaviors as well as those underlying the modifications caused by the feedback are becoming increasingly understood, even on a molecular level. It is straightforward to hypothesize that the constant interaction between ongoing neural activity and the incoming sensory stream allows the organism to balance behavioral flexibility with efficiency to accomplish adaptive behavioral choice in an often unpredictably changing environment. The centrality of this trade-off is reflected in various fields discussing different aspects of this task. Evolutionary biologists study factors specifying whether flexible generalists or efficient specialists will prevail, psychologists search for transitions between flexible goal-directed actions and efficient habits, ecologists study at what point animals cease to efficiently exploit a resource to invest in costly exploration, computational neuroscientists unravel interactions between flexible model-based and efficient model-free learning processes. Animals and humans have evolved over millions of years to become experts in mastering this fundamental trade-off. Focusing on only one aspect of it fails to capture essential processes controlling behavior.

